# Hippocampal representations of possible futures predict motor output in the superior colliculus

**DOI:** 10.64898/2026.06.12.731955

**Authors:** Cameron Wilhite, Loren M. Frank, Massimo Scanziani

## Abstract

The ability to represent possible futures allows animals to evaluate different options and act accordingly. The hippocampus can express representations of possible future paths, but the brain structures involved in transforming these representations into actions remain unknown. Here we show that hippocampal representations of possible future paths predict motor-related activity in the superior colliculus (SC), a midbrain structure involved in turning. By recording simultaneously from hippocampal neurons and from left-or right-preferring “turn cells” in the motor layers of the SC in navigating mice, we found that hippocampal representations of possible left or right future paths predicted increased firing rates of SC turn cells with matching left or right turn preference. This relationship was also evident in the animals’ behavior: when the hippocampus represented future paths that were not ultimately taken, movement trajectories were nonetheless biased toward those paths, as if animals partially enacted possible futures that were represented but not selected. These results reveal a coordination between a cognitive and a motor structure and provide a potential pathway for mental simulations of possible futures to influence actions.

## Introduction

Animals navigating their environments can mentally represent possible future paths^1–3^. While the expression of these representations allows animals to evaluate multiple options and act accordingly^3–6^, the neural circuits involved in transforming these representations into actions remain unknown.

The hippocampus is capable of representing hypothetical scenarios, including possible future options^1–3^. During locomotion, individual cycles of the ∼8 Hz theta rhythm can include non-local spatial representations that sweep ahead of the animal’s current location toward possible future locations, including along possible future paths^1,2,7^. Whether the expression of these representations leaves a specific and measurable impression on activity in regions that drive action is not known.

To address this, we recorded simultaneously from the hippocampus and the superior colliculus (SC), a conserved midbrain structure involved in turning behavior^8,9^, as mice navigated a Y-maze. We recorded from the intermediate and deep layers of the SC (dSC), layers that contain motor command neurons for left or right turns^10–14^. Furthermore, the dSC is known to integrate diverse inputs related to turning^15–17^, raising the possibility that hippocampal representations of possible futures are integrated at the level of the dSC to influence turning actions. Such a mechanism would imply a consistent, predictive relationship between hippocampal representations of left or right future paths and the activity of dSC neurons associated with left or right turns.

To investigate this relationship, we took advantage of the fact that, as animals approach a bifurcation on a maze, the hippocampus can generate representations that sweep either toward the path the animal will ultimately take or toward the alternative path. Thus, for a given path taken by the animal, we can determine whether the activity of dSC neurons varies depending on the direction of hippocampal sweeps. We show that hippocampal representations of possible future paths predict increased firing rates of dSC neurons with matching turn preference. Moreover, we show that this relationship manifests in behavior: hippocampal representations of future paths are associated with deviations in the animal’s trajectory toward the represented path. These results reveal a coordination between cognitive representations of possible future paths and motor commands that steer the animal toward those paths, suggesting a pathway for mental simulations of possible futures to influence actions.

## Results

### Simultaneous monitoring of CA1 and dSC activity during the Y-maze task

We used silicon probes to record from both hippocampal CA1 and dSC neurons as mice performed a Y-maze spatial navigation task (Figure 1A, Figure S1). Each task session consisted of multiple trial blocks of 50-80 trials during which two of the three reward ports (located at the ends of the maze arms) were active, with their locations changing uncued across blocks. This trial block design was chosen to maintain cognitive engagement with the task. During each trial, the mouse navigated from one reward port to the other and, across trials, consistently shuttled between the two active ports, resulting in turns at the maze bifurcation.

**Figure 1:**
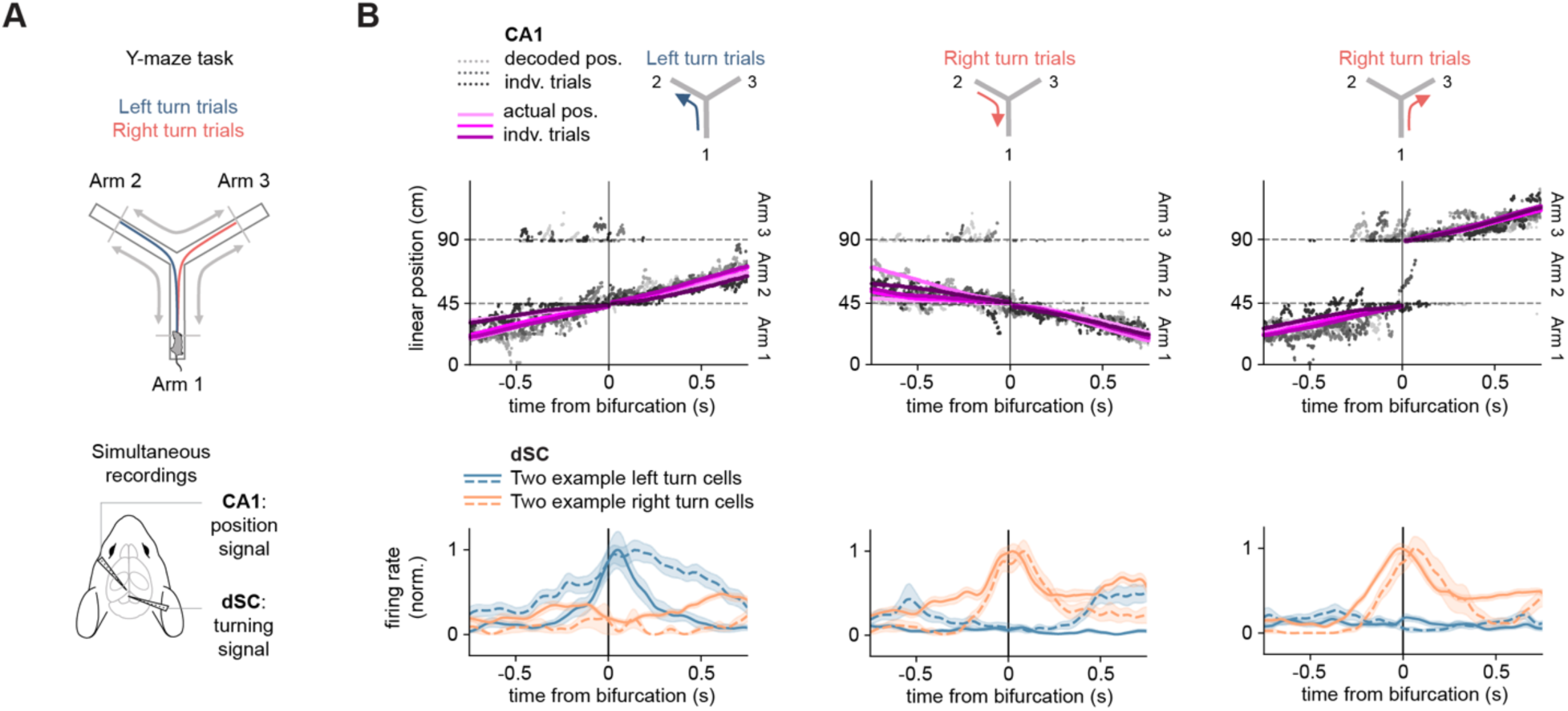
Simultaneous monitoring of hippocampal spatial representations and dSC turn cell activity during a Y-maze task **(A)** Top, Y-maze with average behavioral trajectories of left and right turns across all sessions and mice overlaid (n = 19 sessions, 5 mice). The crossing of the 70% point on the maze arms (gray lines) determines the start and stop of each trial. Bottom, recording configuration targeting both the dorsal CA1 region of the hippocampus and the intermediate and deep layers of the SC (dSC). **(B)** Top, example sets of individual maze traversals (5 trials each, overlaid) as the animal approached the bifurcation from three different maze arms (left panel: left turn trials from arm 1 to arm 2, middle panel: right turn trials from arm 2 to arm 1, right panel: right turn trials from arm 1 to arm 3). Gray scatter points: decoded position from CA1 taken as the peak of the probability distribution at each time point (y-axis, linearized position). Pink lines: the animal’s actual position. Line shading is matched across decoded and actual positions within each trial (i.e., lightest gray trial corresponds to lightest pink trial). Note that, prior to the bifurcation, the decoded position periodically sweeps ahead, representing both the path ultimately taken by the animal and, on some trials, the alternative path. Bottom, two example left turn cells and two example right turn cells recorded simultaneously from the dSC. Solid and dashed lines indicate the average firing rates of two individual cells of the same turn preference (blue: left turn cells, orange: right turn cells) while shaded bands represent s.e.m. (left panel: all left turn trials from arm 1 to arm 2, middle panel: all right turn trials from arm 2 to arm 1, right panel: all right turn trials from arm 1 to arm 3). Note how each turn cell fires similarly at the maze bifurcation regardless of which maze arm the animal approaches the bifurcation from.

We first characterized the CA1 representation of position as animals traversed the Y-maze. We used a state-space decoding algorithm^18^ to estimate the locations represented in the firing patterns of hippocampal neurons (1 ms bins, n = 2642 hippocampal CA1 neurons across 5 animals). We confirmed accurate decoding of position (absolute error = 3.8 ± 0.2 cm, average ± s.e.m.; here and throughout unless stated otherwise). Furthermore, as animals approached the bifurcation, we observed non-local spatial representations that swept left or right ahead of the animal, reflecting possible future paths along the left or right arms of the maze (Figure 1B, top panels).

Next, we focused on “turn cells” in the dSC, a population of neurons that exhibited turn-direction selective firing around the maze bifurcation. We defined turn cells as neurons that showed significant firing rate modulation around turns (significant Turn Direction Selectivity Index or TDSI, see Methods), consistently in the same direction (i.e., same sign TDSI, negative TDSI: left turn-selective, positive TDSI: right turn-selective) regardless of which maze arm the animal came from. As observed in our companion paper^19^, turn cells accounted for ∼30% of dSC cells (167/552 from 5 animals, Figure S2A). Turn cell firing peaked around the bifurcation (+34 ± 172 ms, average ± s.d., Figure S2C, bottom panel) and became directionally selective ∼250 ms before the bifurcation (−248 ± 216 ms, average ± s.d., Figure S2C, top panel). A subset of turn cells (25%, 41/167) became directionally selective prior to the time at which the animals’ behavior diverged between left and right turns (Figure S2B and S2C) in line with premotor activity previously reported in the dSC^12,13,20–23^. Furthermore, consistent with the lateralization of the dSC hemispheres for left and right turns^12,13,20,21^, about two thirds of turn cells showed selective firing for contraversive turns relative to the recorded SC hemisphere (Figure S2D, contraversive: 72%, 120/167, ipsiversive: 28%, 47/167). Together, these observations are consistent with these cells functioning as motor command neurons for turning.

### Turn cell activity is modulated according to hippocampal representations of future paths

Is turn cell activity modulated according to hippocampal representations of future paths? To address this question, we identified periods during which, as the animal approached the bifurcation, the hippocampal representation of position swept ahead, representing future paths along the left or right arm (Figure 2A and 2B, see Methods).

**Figure 2:**
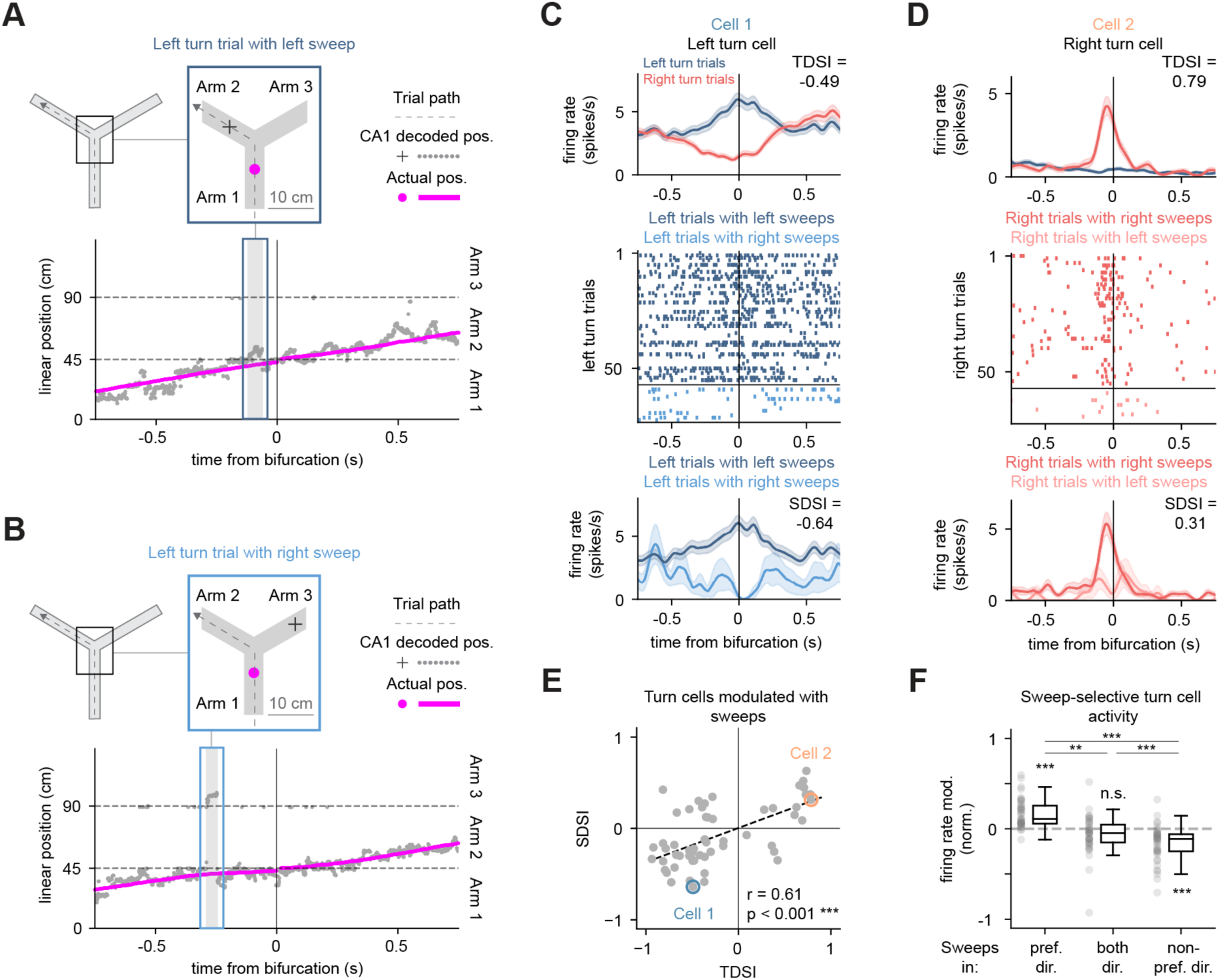
Turn cell activity is modulated in coordination with hippocampal sweeps. **(A)** Example left turn trial with left hippocampal sweep. Top (expanded view), maze schematic showing a snapshot of the animal’s actual position (pink circle) and decoded position (black crosshair) during the sweep. The position of each marker reflects the average linearized position, either actual or decoded, across the time window indicated by the outlined gray band in the bottom plot. Bottom, plot showing the decoded position (gray) and actual position (pink) relative to the maze bifurcation (y-axis, linearized position). Note the decoded position sweeping up into the left arm as the animal approaches the bifurcation. **(B)** Same as in A, but for a left turn trial with a right hippocampal sweep during the same session. **(C)** Top, firing rate profiles of a left turn cell in the dSC (Cell 1, negative TDSI). Data taken from the same session as in A and B. Middle, spike raster plot for Cell 1 during left turn trials. Dark blue spikes indicate spikes during left turn trials with left sweeps; light blue spikes indicate spikes during left turn trials with right sweeps. Bottom, firing rate profiles of Cell 1 during left turn trials separated by sweep direction (same color scheme as in middle panel). Note increased firing during left turn trials with left sweeps compared to left turn trials with right sweeps corresponding to a significantly negative Sweep Direction Selectivity Index value (SDSI: p < 0.01, permutation test). **(D)** Same as in C but for a right turn cell (Cell 2, positive TDSI) during right turn trials. Data taken from a different session than in A-C. Note increased firing during right turn trials with right sweeps compared to right turn trials with left sweeps corresponding to a significantly positive SDSI value (p < 0.05, permutation test). **(E)** Correlation between TDSI and SDSI across cells whose activity was significantly modulated with sweeps (n = 56 cells, Pearson r = 0.61, p < 0.001). “Sweep-selective turn cells” are those in the lower-left and upper-right quadrants. **(F)** Firing rate modulation of sweep-selective turn cells during trials with sweeps only in the cell’s preferred direction, non-preferred direction, or both directions, normalized by the cell’s firing during trials without sweeps. Note bidirectional modulation of firing rates compared to the no sweep condition (preferred/non-preferred only: p < 0.001, both: p > 0.05, 1-sample t-tests, Bonferroni-corrected). These distributions were significantly different from one another (pref. vs both: p < 0.01, pref. vs non-pref: p < 0.001, and both vs non-pref: p < 0.001, paired t-tests, Bonferroni-corrected).

As expected^24,25^, most hippocampal sweeps were directed toward the path the animal would ultimately take (“ipsi sweeps”, 73.7 ± 1.8% across sessions) and the rest toward the alternative path (“contra sweeps”). Contra sweeps occurred earlier relative to the turn and swept farther ahead of the animal compared to ipsi sweeps (Figure S3A and S3B). Moreover, while the onsets of both ipsi and contra sweeps occurred preferentially during late phases of the ∼8.5 Hz (8.49 ± 0.07 Hz) hippocampal theta rhythm, contra sweeps exhibited weaker phase-locking compared to ipsi sweeps (Figure S3C). Lastly, consistent with recent results in rats^25^, the expression of contra sweeps was transiently increased following trial block switches compared to ipsi sweeps (Figure S3D).

To determine whether, for a given turn direction taken by the animal, dSC turn cell activity varied depending on the direction of hippocampal sweeps, we focused on trials with only right or left sweeps, excluding trials with sweeps in both directions and trials without sweeps. For each cell, we computed a Sweep Direction Selectivity Index (SDSI, see Methods) that quantified, for turns in a given direction, the difference in firing rates seen on trials with right sweeps versus trials with left sweeps, following the same sign convention as the TDSI (negative SDSI: left sweep-selective, positive SDSI: right sweep-selective).

Strikingly, the activity of a third of turn cells was significantly modulated around the time of the bifurcation depending on the direction of hippocampal sweeps (34%, 56/167 cells, see Methods for inclusion criteria, Figure 2C, cell 1, SDSI: p < 0.01, Figure 2D, cell 2, SDSI: p < 0.05, permutation tests). Moreover, there was a strong positive correlation between TDSI and SDSI across this population (Figure 2E, n = 56 cells, Pearson r = 0.61, p < 0.001; Figure S5A, distribution across animals). Indeed, most of these cells (75%, 42/56) fired more on average during trials with sweeps in their preferred turn direction compared to trials with sweeps in their non-preferred turn direction (“sweep-selective turn cells,” lower-left and upper-right quadrants of Figure 2E). Of these, most cells exhibited sweep-direction selective firing during trials in which the animal turned in the cell’s preferred turn direction (62%, 26/42, Figure 2C, 2D, S4B and S4C), while the rest exhibited selectivity during trials in which the animal turned in the cell’s non-preferred turn direction (Figure S4D).

By selecting the cells with a significant SDSI, we thus observed a strong positive correlation between TDSI and SDSI (Figure 2E). Does this correlation extend to the population of cells whose SDSI did not reach the threshold for significance? Across this population (n = 98 cells, see Methods for inclusion criteria), there remained a significant correlation between TDSI and SDSI. Specifically, this correlation was positive during trials in which the animal turned in the cell’s preferred turn direction (Figure S5B, left panel, TDSI vs. SDSI, n = 98 cells, Pearson r = 0.44, p < 0.001) and was absent during trials in which the animal turned in the cell’s non-preferred turn direction (Figure S5B, right panel, Pearson r = −0.16, p > 0.05). These results indicate that, rather than being a feature of a discrete subset of neurons, sweep-related modulation of turn cell activity is a continuum across the turn cell population.

Lastly, to determine whether the activity of turn cells can be bidirectionally modulated, that is increase or decrease their firing in coordination with sweep direction, we analyzed the firing rates of each sweep-selective turn cell (n = 42 cells) around the time of the bifurcation across three distinct trial types: trials with sweeps 1) only in the cell’s preferred direction, 2) only in the cell’s non-preferred direction, and 3) in both directions, all relative to trials without sweeps. This analysis revealed a clear bidirectional modulation of dSC firing rates relative to the no-sweep condition: sweep-selective turn cells fired more during trials with sweeps only in their preferred direction, less during trials with sweeps only in their non-preferred direction, and showed no significant change during trials with sweeps in both directions (Figure 2F, preferred/non-preferred only: p < 0.001, both: p > 0.05, 1-sample t-tests). Together, these results reveal that the activity of dSC turn cells, both at the single-cell level and at the population level, is modulated in coordination with hippocampal representations of future paths.

### Hippocampal representations of future paths reliably predict subsequent dSC activity

Hippocampal sweeps can occur at various time points prior to the bifurcation. How early before the bifurcation do sweeps predict the activity of dSC cells at the bifurcation? To address this question, we used generalized linear models (GLMs, see Methods). As sweeps typically occurred in-phase with the hippocampal theta rhythm (Figure S3C), we divided each trial into theta cycles prior to and at the time of the bifurcation (cycles −4 through −1 and cycle 0, respectively) and, for each of these cycles, assessed whether the presence and direction of sweeps predicted dSC firing at the bifurcation (Figure 3A).

**Figure 3:**
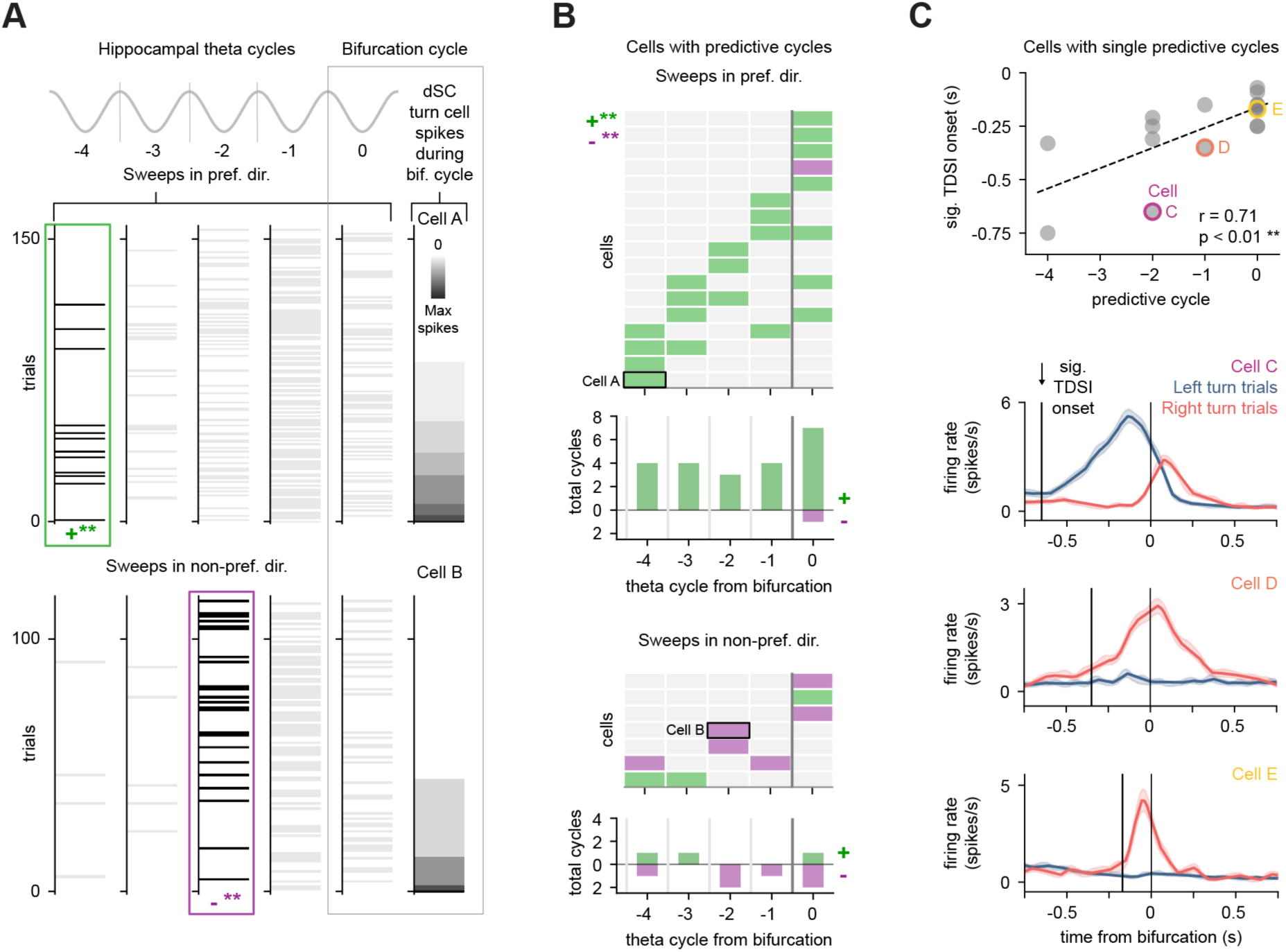
Hippocampal sweeps reliably predict future turn cell activity. (A) GLM analysis to assess whether the presence and direction of hippocampal sweeps during individual theta cycles prior to and at the time of the bifurcation (cycles −4 through −1 and cycle 0, respectively) predicts dSC turn cell firing at the bifurcation. Top, GLM analysis for an example dSC turn cell (Cell A). Spike count for Cell A during the bifurcation cycle (cycle 0) is shown at the far right. Trials are sorted by increasing spike counts of Cell A. Hippocampal sweeps in Cell A’s preferred turn direction across trials are plotted as horizontal lines during their corresponding theta cycle relative to the bifurcation (cycles −4 through 0). Trials are those in which the animal turns in the cell’s preferred turn direction. Note that sweeps occurring during cycle −4 are associated with increases in spiking of Cell A during the bifurcation cycle. Significantly positive (+**) predictive cycle is shown by a green outline (p < 0.01, Wald z-tests). Bottom, GLM analysis for another example turn cell (Cell B) in which sweeps in Cell B’s non-preferred turn direction are shown. Note that sweeps occurring during cycle −2 across trials are associated with decreases in spiking of Cell B during the bifurcation cycle. Significantly negative (−**) predictive cycle is shown by a purple outline (p < 0.01, Wald z-tests). Sweeps occurring during significantly predictive cycles are shown in black to highlight the relationship between sweeps and turn cell firing. Sweeps occurring during non-significantly predictive cycles are shown in gray. (B) Top, all cells with predictive cycles for sweeps in the cell’s preferred turn direction sorted by theta cycle from bifurcation. Bottom, all cells with predictive cycles for sweeps in the cell’s non-preferred turn direction. (C) Top, correlation between timing of predictive cycle and onset of significant TDSI across turn cells. This analysis was necessarily done only on cells whose activity was either positively or negatively predicted by sweeps during a single theta cycle (n = 16 cells, Pearson r = 0.71, p < 0.01). Bottom, firing rate profiles of three example turn cells (Cells C, D, and E from top plot) showing progressively later significant TDSI onsets relative to the turn.

Strikingly, sweeps occurring up to 4 cycles (∼500 ms) before the bifurcation could significantly predict dSC activity at the bifurcation (Figure 3B, p < 0.01, Wald z-tests). “Predictive cycles” (i.e., theta cycles during which sweeps significantly predicted changes in dSC activity at the bifurcation) were observed for 50% of sweep-selective turn cells and for 7.6% of all possible cycles (Figure 3B, cells: 21/42, cycles: 32/420), both of which significantly exceeded shuffled controls (Figure S6A, shuffled data: cells, 29 ± 0.2%, p = 0.001; cycles, 4.1 ± 0.03%, p < 0.001, permutation tests). Most predictive cycles were driven by sweeps in a cell’s preferred turn direction (72%, 23/32). Of these, most cycles predicted increases in activity (Figure 3B, top panels, 96%, 22/23), whereas, for predictive cycles driven by sweeps in a cell’s non-preferred direction (28%, 9/32) most cycles predicted decreases in activity (Figure 3B, bottom panels, 67%, 6/9 cycles).

Distinct dSC turn cells become directionally selective at different times prior to the bifurcation (−248 ± 216 ms, average ± s.d., Figure S2C, top panel). This raises the possibility that those dSC turn cells that showed earlier directional selectivity would also be those whose firing, at the bifurcation, is modulated in coordination with early sweeps. This was indeed the case: focusing on cells with a single predictive cycle, we found a positive correlation between the time of predictive cycles and the time at which turn cells became directionally selective (Figure 3C, top panel, n = 16 cells, Pearson r = 0.71, p < 0.01). In other words, sweeps occurring during earlier theta cycles predicted the activity of turn cells that became directionally selective earlier, and vice versa. Thus, hippocampal sweeps can predict turn-related activity in the dSC, and the timing of this predictability is related to the time at which individual turn cells become directionally selective.

Given that hippocampal sweeps can predict dSC activity at the bifurcation, we next asked whether an interaction in the opposite direction also occurred. That is, can dSC activity prior to the bifurcation predict hippocampal sweeps at the bifurcation? This possibility would be consistent with a direct influence of head turning behavior on hippocampal representations^5^. To address this question, we again turned to GLMs (see Methods). Mirroring the GLM analysis above (Figure 3), we divided each trial into theta cycles prior to and at the time of the bifurcation and, for each of these cycles, assessed whether dSC activity in individual turn cells predicted the presence and direction of hippocampal sweeps at the bifurcation.

We identified only a single instance of a significant cycle in which dSC turn cell activity predicted hippocampal sweeps at the bifurcation (Figure S7A, p < 0.01, Wald z-test). Additional analyses using a less conservative significance threshold (p < 0.05, Wald z-tests) yielded only weak effects that did not robustly exceed shuffled controls (Figure S7B and S7C). Thus, while dSC activity does not reliably predict hippocampal sweeps at the bifurcation, hippocampal sweeps reliably predict dSC activity.

### Hippocampal sweep direction matches deviations in behavioral trajectory

dSC activity impacts turning behavior. Given that the direction of hippocampal representations of future paths predicts differences in dSC activity, we asked whether it also predicts differences in the animal’s behavior. This was indeed the case.

First, for a given turn direction, turns were tighter during ipsi sweep trials compared to contra sweep trials (Figure 4A). Trajectories during contra sweep trials deviated more from a linear path and exhibited greater cumulative angular change (IdPhi^24^) compared to ipsi sweep trials (Figure 4B and 4C, see Methods). Second, we observed an effect of sweep presence and direction on trial duration. Trials without sweeps had the shortest duration (median: 1.73 s), followed by ipsi sweep trials (1.83 s), contra sweep trials (1.95 s), and trials with sweeps in both directions (2.22 s; Figure S8A). Consistent with this effect, running speed was highest on trials without sweeps (median: 34.1 cm/s), followed by ipsi sweep trials (32.1 cm/s), contra sweep trials (31.2 cm/s), and trials with sweeps in both directions (27.5 cm/s; Figure S8B). Thus, the movement trajectory of the animal around the turn and its running speed reflect the presence and direction of hippocampal sweeps.

**Figure 4:**
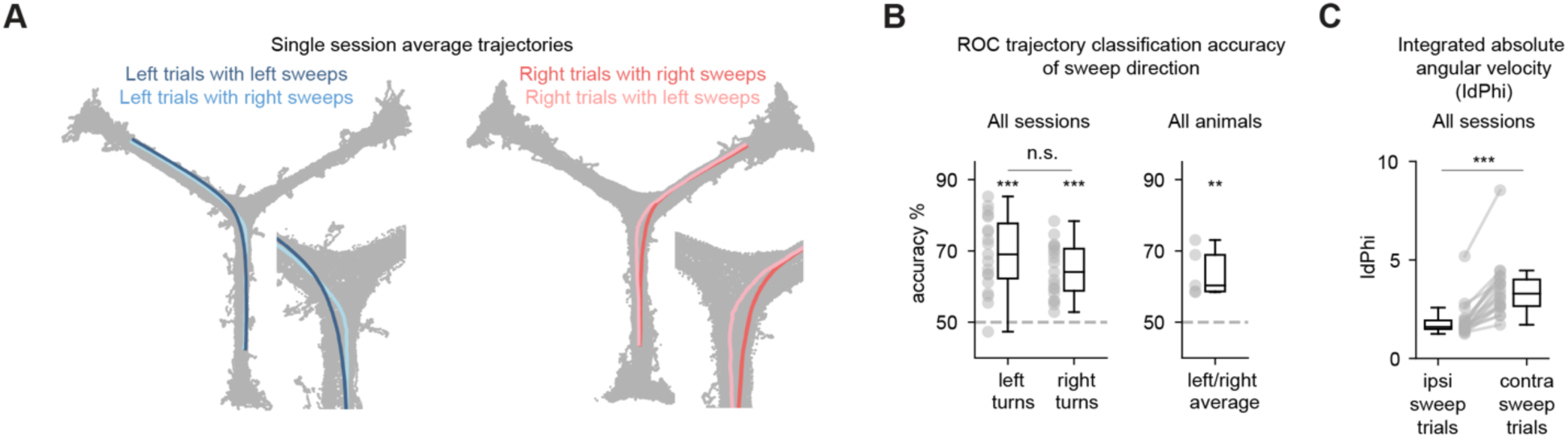
Hippocampal sweep direction and deviations in trajectory. (A) Left, average left turn trajectories for a single session color coded by sweep direction. Dark blue trajectory indicates average of left turn trials with left sweeps (ipsi sweep trials), light blue trajectory indicates average of left turn trials with right sweeps (contra sweep trials). Session is the same session as in Figure 2A, 2B, and 2C. Note deviation of trajectory for contra sweep trials toward the alternative path. Right, same as in left panel but for right turn trials. Session is the same session as in Figure 2D. (B) Left, ROC classification accuracy of sweep direction from deviation in behavioral trajectory from a linear path around the time of turns (see Methods), shown across sessions for both turn directions (n = 19 sessions, left turns: 69 ± 2%, right turns: 64 ± 2%, left and right turns: p < 0.001, 1-sample t-tests compared to 50%, Bonferroni-corrected; left vs. right turns: p > 0.05, paired t-test). Right, ROC classification accuracy between ipsi and contra sweep trials, combined across left and right turns per animal (n = 5 animals, 64 ± 3%, p < 0.01, 1-sample t-test compared to 50%). (C) Integrated absolute angular velocity (IdPhi) values calculated across the average trajectories for ipsi vs. contra sweep trials for each session (n = 19 sessions, p < 0.001, paired t-test). Values are averaged between left and right turns.

## Discussion

We identified a consistent, predictive relationship between representations of possible future paths in the hippocampus and the activity of dSC neurons associated with turning behavior (i.e., turn cells). Furthermore, consistent with the role of dSC activity in generating turns, when the hippocampus represented possible paths that were not ultimately taken, movement trajectories were nonetheless biased toward those paths, as if animals partially enacted possible futures that were represented but not selected.

In our companion paper^19^, we show that dSC turn cells can also be modulated in coordination with the stepping rhythm. Neurons modulated in coordination with the stepping rhythm and neurons modulated in coordination with hippocampal sweeps are not separate populations. In fact, approximately half of the dSC turn cells whose activity was modulated in coordination with hippocampal sweeps exhibited activity that was significantly phase-locked to the stepping cycle (48%, 27/56 cells).

While our study reveals a unidirectional predictive relationship between hippocampal sweeps and dSC turn cell activity, the data are correlational and do not establish a causal role for hippocampal sweeps in modulating dSC turning commands. Furthermore, a recent study in mice has shown that SC activity is causally linked to changes in virtual heading during REM sleep^12^, suggesting that dSC activity can shift internal representations of space. Thus, we cannot exclude that dSC activity may influence the direction of hippocampal sweeps. Alternatively, common upstream inputs to these structures, or a third sensorimotor or behavioral variable that jointly influences hippocampal and dSC activity could explain the results presented here.

Nevertheless, given our results that hippocampal sweeps predict future turn cell activity and that this relationship correlates with deviations in the animals’ trajectory toward the represented path, we favor the possibility that hippocampal sweeps may modulate dSC motor commands to bias turning behavior. How might such a modulation take place? Our results provide support for an integration-like mechanism in the dSC related to representations of possible futures. Previous work has demonstrated that integration is a key function of the dSC, wherein inputs related to orienting decisions drive progressive increases in activity that, upon reaching a threshold, drive action^26–29^. Our GLM analysis shows that hippocampal sweeps occurring up to ∼500 ms before the maze bifurcation can predict dSC activity at the bifurcation (Figure 3A and 3B). This timing suggests that hippocampal sweeps may be transmitted to the dSC to modulate turning commands, perhaps in a sequential manner aligned with the time at which individual turn cells become directionally selective (Figure 3C), to bias turns toward behaviorally relevant locations in nearby space^30–32^. Such a possibility is consistent with several motor-based theories of cognition^33–36^ and with the century-old proposal that the mere idea of a movement automatically triggers the movement itself^37–39^. However, further manipulation experiments are necessary to establish a causal role for hippocampal sweeps in modulating dSC activity and the anatomical pathways involved in this process.

Cognitive abilities such as imagination, planning, and decision-making involve mentally exploring possible options^2,3^. To guide behavior, these mental explorations must, at some level, influence neural circuits that generate action. Here we show that when the brain simulates a possible future path, it also engages motor commands that actively turn the animal toward that path. These results reveal a coordination between a cognitive and a motor structure and offer a potential mechanism for mental simulations of possible futures to influence actions.

## Acknowledgements.

We thank P. Saraf, L. Ruan and Q.Y. Wu for technical support and the members of the Scanziani laboratory for helpful discussions of this project. Claude (by Anthropic) was used to draft portions of the methods and figure legends, with the authors’ notes and code used as input. This project was supported by the Neuroscience Training Grant T32 (C.W.), by the Discovery Fellowship of the UCSF Graduate Division (C.W.) and the Howard Hughes Medical Institute (M.S. and L.M.F.).

## Author contributions

C.W., L.M.F. and M.S. designed the study. C.W. conducted all experiments and experimental data analysis. C.W., L.M.F. and M.S. wrote the manuscript.

## Methods

### Experimental model and animals

Neural activity was recorded from five male C57BL/6J mice (Jackson Laboratory #000664, 5-6 months old). All mice were housed on a reversed cycle (light/dark cycle 12/12h) with littermates before experimentation and singly housed in enriched cages during food restriction and experimentation. All procedures were conducted in accordance with the regulations of the University of California San Francisco Institutional Animal Care and Use Committee (IACUC).

### Behavioral training and electrode implantation

Implantations were performed in two phases on two distinct days. First, the implantation of the headplate, and second, the implantation of the silicon probes. Mice were implanted under 2% isoflurane anesthesia. The headplate was placed on the skull with dental cement (Unifast LC, GC America; Optibond Universal, Kerr Dental). After post-operative recovery of five days, mice were deprived of food to 85% of their baseline weight and pretrained to run on a 1 m linear track for liquid food reward (sweetened soymilk). After pretraining for one to three days on the linear track, mice were pretrained on the Y-maze task for three to five days (See Y-maze task and trial segmentation) and implanted with silicon probes. Silicon probes were mounted on 3D-printed microdrives to record unilaterally from the superior colliculus (SC) and from the CA1 region of the hippocampus in the opposite hemisphere. The SC data presented here were also used in our companion paper^19^. The types of probes used included: Single-shank (Diagnostic Biochips (DBC) P64-4), four-shank (DBC P64-1, DBC P128-6), and eight-shank probes (DBC P128-8). Probe shanks were inserted through craniotomies at the coordinates described in the Electrode insertion section below. The shanks were initially lowered to 0.7-1.2 mm (SC) and 0.6 mm (CA1) below brain surface and the microdrive was then fixed to the headplate. A ground electrode (0.005” diameter stainless wire, A-M systems) was inserted in the cerebellum. After post-operative recovery, mice were placed back on the Y-maze training protocol for 1-5 days while the shanks were lowered to the intermediate and deep SC (dSC) and to CA1 to ensure that peak Y-maze performance coincided with the time at which the recording sites reached their targets. Mice remained on food restriction for the remainder of the experiment.

### Electrophysiological recordings and video tracking

The data presented in this paper are from three to four 60–140 min sessions per mouse on the Y-maze task. Electrophysiological data were acquired using an Intan RHD2000 system (Intan Technologies LLC) band-pass filtered between 0.1 Hz and 7.5 kHz and digitized at 20 kHz. Spike sorting was performed semi-automatically, using Kilosort 2.0 and Phy^40^. Video data was acquired at 60 frames per second using a CMOS camera (Basler acA1300-200um) placed above the maze. A machine-learning algorithm, DeepLabCut^41^, was trained to track features of the probe housing and distinct body parts of the mice (microdrive probe base (head), left and right ears, base of the tail). For position analysis and trial segmentation on the Y-maze, the probe base position (head position) was used as the actual position of the mouse.

### Y-maze task and trial segmentation

Mice navigated an elevated Y-maze (arm length: 45 cm, arm width: 5 cm) in search of liquid food reward, which was dispensed from two out of three ports at the ends of the maze arms. The task consisted of six to eight 50–80 trial blocks, with the locations of the active ports switching in an uncued manner between blocks. The active ports were chosen randomly within each block. Animals could not visit the same port on consecutive trials to receive a reward, so mice learned to alternate between the active ports. For trial segmentation, we used the track_linearization package^42^ to map the animal’s position on the Y-maze to a one-dimensional representation with a corresponding maze arm identifier. We then defined a position threshold at 70% of the length of each maze arm (31.5 cm measured outward from the center of the maze or 13.5 cm measured inward from the end of the arm). Trials were defined as periods from when the animal, after leaving a reward port, crossed the position threshold of one arm, made a turn, and crossed the position threshold of another arm without entering any other arms. The time at which an animal reached the bifurcation was defined as the time at which the animal’s linearized head position transitioned from one arm to another at the center of the maze. Animals took ∼1.8 seconds to complete each trial (median: 1.85 s, IQR: 1.58–2.23 s, n = 8651 trials, 5 mice).

### Head angle relative to the track and time of behavioral divergence

Head angle relative to the track was defined as the angle between the animal’s head direction vector (given by the vector perpendicular to the line connecting the ears) and the midline of the track segment from which the animal approached the bifurcation. The time of behavioral divergence was defined as the time at which the animal’s head angle relative to the track significantly diverged between left and right turns. To determine the average time of behavioral divergence across sessions and animals (n = 19 sessions, 5 animals, Figure S2B), we calculated a Behavioral Divergence Index (BDI) as the difference between session-averaged head angles relative to the track for left and right turn trials. BDI was calculated within 200 ms windows advanced in 20 ms steps from 0.85 s before to 0.85 s after the time at which the animal reached the bifurcation. BDI significance was assessed using a permutation test: for each window, a null distribution was generated by randomly permuting left and right turn labels across sessions (10,000 permutations) and recalculating BDI from the resulting averages. Windows were considered significant only when the observed BDI was more extreme than all shuffled values (p < 1/10,000). The time of behavioral divergence was defined as the earliest time window belonging to a continuous sequence of such significant windows.

### Electrode insertion, electrode coordinates, histology and recording site assessment

In three mice, the right SC was targeted; and in two mice, the left SC was targeted. Lambda was used as the skull reference for SC craniotomies, while bregma was used for hippocampal craniotomies. The SC was targeted by a craniotomy at AP: 0 mm from lambda, ML: ± 2.5 mm, and the silicon probe was inserted at a 32° angle from vertical along the anterior-posterior axis of the SC. The CA1 region of the hippocampus was targeted in the opposite hemisphere by a craniotomy at AP: −2.0 mm from bregma, ML: ± 1.5 mm, and the silicon probe was inserted vertically, with the shank plane rotated 45° from the sagittal plane along the long axis of the hippocampus. Prior to implantation, the backs of all probe shanks were coated with DiI solution (Fisher, V22885) for probe track localization using a fluorescence microscope (Figure S1A). The shanks were initially lowered to 0.7–1.2 mm (SC) and 0.6 mm (CA1) below the surface of the brain (dorsoventral, DV) and advanced over consecutive days to reach the dSC and cell body layer of CA1. The dSC was targeted using DV coordinates, and CA1 was targeted using DV coordinates and physiological signatures such as sharp-wave ripples. Once the target regions were reached, dSC probes were advanced between sessions (days) to record from different neurons while CA1 probes were slightly advanced between sessions to maintain maximal units for decoding. For the dSC, the average depth for the first recording was DV: −1.51 ± 0.13 mm measured from the topmost channel of the probe while the average depth for the last recording was DV: −2.96 ± 0.06 mm measured from the bottommost channel of the probe (Figure S1B, left panel). For CA1, average depths were DV: −1.00 ± 0.11 mm and DV: −1.39 ± 0.10 mm, respectively (Figure S1B, right panel).

For anatomical analysis, mice were perfused transcardially with phosphate-buffered saline (PBS) and then with 4% paraformaldehyde (PFA) in PBS. Brains were extracted, stored in 4% PFA overnight at 4°C and cut with a vibratome to 100 µm thick coronal sections. Slices were mounted in DAPI Vectashield (Vector Laboratories H1500), and fluorescence images were acquired with an Olympus MVX10 MacroView microscope. Probe tracks were identified using the DiI signal.

### Turn Direction Selectivity Index (TDSI) and turn cells

Turn-direction selectivity was quantified using a Turn Direction Selectivity Index (TDSI), calculated as the difference between the average firing rates during right and left turns divided by their sum ((Right FR - Left FR) / (Right FR + Left FR)) over a 200 ms window centered on the time of the cell’s peak firing rate. Thus, right turn cells were defined by positive values and left turn cells by negative values. To obtain firing rates, spike trains were converted to binary vectors, smoothed with a Gaussian kernel (50 ms), normalized by the kernel width, and averaged across trials. To determine the time window for the TDSI calculation, we first defined a wide window of ± 0.5 s around the time at which the animal reached the bifurcation and, for each neuron, identified within this window the time of peak firing rate for either left or right turns (whichever was greater) using the neuron’s average firing rate across all arms. TDSI was then computed across each arm within a narrow window of ± 0.1 s centered on this peak firing time.

TDSI significance was assessed using a permutation test. For each neuron and maze arm, left and right turn trials were pooled together and randomly reassigned into two groups whose sizes matched the original sample sizes. TDSI was then recomputed within the narrow window centered on the same actual peak firing time. This procedure was repeated 1000 times to generate a shuffled distribution of TDSI values. For right turn cells, the p-value was defined as the fraction of shuffled TDSI values that were greater than or equal to the observed TDSI, or less than or equal to the observed TDSI for left turn cells. Turn cells were defined as neurons whose TDSI retained the same sign and remained significant (p < 0.05) across all maze arms.

### Instantaneous TDSI and significant TDSI onset

We computed instantaneous TDSI across the entire trial period^10^ to determine the time at which turn cells became directionally selective (significant TDSI onset, Figure S2A). TDSI was calculated within 200 ms windows advanced in 20 ms steps from 0.85 s before to 0.85 s after the time at which the animal reached the bifurcation. TDSI significance was assessed with a permutation test: for each window, left and right turn labels were randomly permuted across trials and TDSI was recomputed. This process was repeated 500 times to generate a null distribution for each window. Windows were considered significant when the observed TDSI was more extreme than all shuffled values (p < 1/500). Significant TDSI onset was defined as the earliest time point at which a turn cell exhibited two consecutive significant TDSI windows in the expected direction (e.g., for a left turn cell, two consecutive significant TDSI windows with negative TDSI).

### Spatial decoding from hippocampal activity

We used an established state-space decoding algorithm from Denovellis et al., 2021^18^ to decode spatial position from populations of clustered hippocampal neurons. Briefly, for each session we built an encoding model to relate hippocampal spiking to the animal’s linearized head position in 1 ms bins. We then decoded position from hippocampal spiking in 0.5 cm linear position bins and 1 ms temporal bins. For cross-validation, we trained the encoding model on 75% of the data in each session and tested it on the remaining 25%, repeating this across four folds. We used a single-state continuous, causal decoding model^18^ in which the continuous state was modeled by a random walk transition matrix with a 9.6 cm movement variance. To obtain the decoded position, the posterior probability distribution of position was first smoothed across spatial bins (i.e., along the position axis) using a 2.5 cm moving window. The decoded position in each time bin was then defined as the location of the peak of this smoothed posterior. For each time bin, we also classified the animal’s actual and decoded positions into one of three maze arms.

### Identification of hippocampal sweeps from decoded position

Sweep times were identified as times during trial periods (i.e., during locomotion) in which the decoded position was in a maze arm distinct from the animal’s actual position for at least 15 consecutive time bins (15 ms). This duration requirement was used to prevent rapid, spurious jumps in the decoded position from being identified as sweeps. Only sweeps that occurred prior to the bifurcation were analyzed. Sweeps were classified as ipsi sweeps if they proceeded into the maze arm that the animal would eventually take or contra sweeps if they proceeded into the alternative arm. For trial-based analyses, trials were classified as ipsi sweep trials (containing only ipsi sweeps (1 or more)), contra sweep trials (containing only contra sweeps), trials with sweeps in both directions (containing both ipsi and contra sweeps), or trials without sweeps.

To compare the sweeps identified here in mice to those observed in rats, we extracted several key features of ipsi and contra sweeps including their timing relative to the hippocampal theta cycle, how far they swept ahead of the animal, and their density around the time of trial block switches (Figure S3). To analyze the timing of sweeps relative to the hippocampal theta cycle, the local field potential (LFP) from the CA1 cell layer was bandpass filtered between 5 and 11 Hz (Butterworth, second-order). Peaks and troughs of the filtered LFP were identified and the instantaneous phase was obtained using the Hilbert transform. The LFP signal was then segmented into half-cycles (peak-to-trough and trough-to-peak). Sweep times were assigned a phase by identifying the half-cycle in which the sweep onset occurred and linearly interpolating the Hilbert phase within that interval. For plotting (Figure S3C), phase 0 corresponds to the trough of the theta cycle at the CA1 pyramidal cell layer. Phase-locking strength was quantified using the Rayleigh Z metric^19,43^, which uses the formula *Z = nR*^2^, where n is the number of events (sweeps) and R is the mean resultant length. For each session, sweep phases were pooled across left and right turn trials, and the preferred phase (circular mean) and Rayleigh Z were computed from the pooled phases. Sweep length (distance) was calculated as the average difference between the animal’s actual and decoded positions during the duration of the sweep. Mirroring hippocampal sweeps observed in rats, ipsi and contra sweeps occurred preferentially during late phases of theta (0 to π rad., Figure S3C)^2,7,25^ and swept one body length ahead of the animal on average (∼9 cm here, Figure S3B, ∼25 cm in rats^7,25^). Furthermore, consistent with a recent report in rats^25^, we observed a transient increase in the density of contra sweeps relative to ipsi sweeps after trial block switches (Figure S3D). Thus, the hippocampal sweeps reported here in mice exhibit strikingly similar features to those in rats.

### Sweep Direction Selectivity Index (SDSI)

To investigate the relationship between dSC turn cell firing and sweep direction, we took advantage of the fact that, for a given turn direction, some trials contain ipsi sweeps and others contain contra sweeps. Thus, for trials in a given turn direction, we could assess whether the activity of individual dSC turn cells varied depending on the direction of hippocampal sweeps. As this analysis necessarily required trials with only ipsi or contra sweeps, we excluded trials with sweeps in both directions and trials without sweeps.

For each dSC turn cell, we then computed a Sweep Direction Selectivity Index (SDSI) around the bifurcation separately for all left turn trials and all right turn trials. SDSI was calculated as the difference between the average spike count during all trials with right sweeps and the average spike count during all trials with left sweeps, divided by their sum ((Right sweep spike count - Left sweep spike count) / (Right sweep spike count + Left sweep spike count)), where spike counts were taken from a 200 ms window centered on the time that the animal crossed the bifurcation (t = 0). Thus, for a given turn direction, cells that fired more at the bifurcation during trials with right sweeps were defined by positive SDSI values and cells that fired more during trials with left sweeps by negative SDSI values.

SDSI significance was assessed using a permutation test. For each neuron and turn direction, all trials with right sweeps and all trials with left sweeps were pooled together and randomly reassigned into two groups whose sizes matched the original sample sizes. SDSI was then recomputed. This procedure was repeated 1000 times to generate a shuffled distribution of SDSI values. For cells that fired more during trials with right sweeps, the p-value was defined as the fraction of shuffled SDSI values that were greater than or equal to the observed SDSI, or less than or equal to the observed SDSI for cells that fired more during trials with left sweeps. Thus, each cell was assigned an SDSI value and an associated p-value for each turn direction. SDSI values were deemed significant if their associated p-value was less than 0.05. 41% of turn cells (69/167 cells) had a significant SDSI value in one or both turn directions (61 cells in one turn direction only, 8 cells in both turn directions). We excluded two subsets of these cells from further analyses. First, we excluded the eight cells that had significant SDSI values in both turn directions, as no single value could be compared against the cell’s TDSI. Second, we excluded five cells that had significant SDSI values in their non-preferred turn direction that were driven by extremely low spike counts, which forced their SDSI values to the maximum (±1) and produced spuriously significant permutation results. We therefore focused on the remaining 56 cells, as each cell had a single, non-spurious, significant SDSI value that could be directly compared with its TDSI (Figure 2E). Of these cells, “sweep-selective turn cells” were defined as those that fired more during trials with sweeps in their preferred direction compared to trials with sweeps in the non-preferred direction (75% of cells, 42/56, lower-left and upper-right quadrants of Figure 2E). The remaining 98 cells with no significant SDSI in either turn direction constitute the population analyzed in Figure S5B. For these cells, we ran two Pearson correlations: TDSI vs. SDSI computed during trials in the cell’s preferred turn direction (Figure S5B, left panel) and TDSI vs. SDSI computed during trials in the cell’s non-preferred turn direction (Figure S5B, right panel).

### Sweep-selective turn cell activity relative to trials without sweeps

To determine whether the activity of sweep-selective dSC neurons can be bidirectionally modulated, that is increase or decrease their firing in coordination with sweep presence and direction, we analyzed the firing rates of sweep-selective turn cells (n = 42, see above section) around the time of the bifurcation across three distinct trial types: trials with sweeps 1) only in the cell’s preferred direction, 2) only in the cell’s non-preferred direction, and 3) in both directions, all relative to trials without sweeps.

For each sweep-selective turn cell, we calculated a normalized firing rate modulation score for each of the three distinct trial types above as the difference between the average spike count during that trial type and the average spike count during trials without sweeps, divided by their sum ((trial type spike count – no sweep spike count) / (trial type spike count + no sweep spike count)), calculated over the same 200 ms bifurcation window used for SDSI. Accordingly, each sweep-selective turn cell was assigned a modulation score for each trial type (Figure 2F). Positive values indicate that a cell fired more on average during that trial type compared to no sweep trials, negative if it fired less, and zero if there was no change.

### Generalized linear models for assessing if hippocampal sweeps predict turn cell firing

We built Poisson family generalized linear models (GLMs, log link function) to determine whether and when hippocampal sweeps prior to the bifurcation could predict dSC activity at the bifurcation across trials (Figure 3). We included trials with sweeps exclusively in one direction (ipsi or contra) as well as trials without sweeps, excluding trials with sweeps in both directions.

We divided each trial into theta cycles prior to and at the time of the bifurcation (cycles −4 through −1 and cycle 0, respectively) and, for each cycle, extracted three binary predictors across trials: turn direction (left = 1, right = 0), left sweep (1 or 0), and right sweep (1 or 0). For each sweep-selective turn cell (n = 42), the response variable was defined as the spike count across trials during the bifurcation cycle (cycle 0). Importantly, separate GLMs were fit for each cycle (−4 through 0), with predictors drawn from that cycle and the response variable fixed as turn cell spike counts at cycle 0.

For each turn cell, we extracted the beta coefficients for the left and right sweep predictors from each GLM, resulting in 10 beta coefficients per cell (2 sweep directions x 5 theta cycles). The significance of each coefficient was assessed using a Wald z-test. Significant beta coefficients using a conservative threshold (p < 0.01) were converted to their sign (±1), indicating whether sweeps during a given theta cycle significantly predicted increased firing (+1, green-colored cycles in Figure 3) or decreased firing (−1, purple-colored cycles in Figure 3) at the bifurcation cycle. Such cycles were deemed “predictive cycles.”

We performed permutation tests (1000 iterations) to assess whether the observed proportion of turn cells with predictive cycles and the observed proportion of total predictive cycles exceeded chance. For each iteration, spike counts for each turn cell were shuffled separately within left and right turn trials to disrupt sweep-firing relationships. P-values for cells and cycles were calculated as the proportion of shuffles in which the shuffled statistic met or exceeded the observed values. Both the proportion of turn cells with predictive cycles and the proportion of predictive cycles significantly exceeded shuffled controls (Figure S6A).

To assess whether adding sweep predictors improved model performance for each significantly predictive cycle (n = 32 total cycles, Figure 3B), we compared a full model (turn direction predictor + sweep predictors) to a reduced model (turn direction predictors only) using 5-fold cross-validation. Model improvement was quantified as percentage reduction in mean absolute error (MAE) between the reduced model and the full model calculated as: (MAE reduced model - MAE full model) / MAE reduced model × 100. The observed positive values across folds indicate that adding sweep predictors improved model prediction beyond turn direction alone (Figure S6B).

### Generalized linear models for assessing if turn cell firing predicts hippocampal sweeps

We built binomial family GLMs (logit link function) to test whether and when dSC turn cell firing prior to the bifurcation could predict the direction of hippocampal sweeps at the bifurcation across trials (Figure S7). For this analysis, we included trials with sweeps exclusively in one direction (ipsi or contra), excluding trials with sweeps in both directions as well as trials without sweeps.

As in the previous GLM analysis, we divided each trial into theta cycles spanning cycles −4 through 0 from the bifurcation. For each cycle, we extracted a binary turn-direction predictor (left turn = 1, right turn = 0) and spike counts for a given sweep-selective turn cell across trials. For the response variable, we identified whether a hippocampal sweep occurred in cycle 0, and, if so, assigned sweep direction as left = 1 and right = 0. Separate GLMs were fit per cycle (−4 through 0), with predictors drawn from that cycle and the response variable fixed as sweeps at cycle 0.

For each sweep-selective turn cell (n = 42), we extracted the beta coefficients for the spike-count predictors from cycles −4 through 0 in each GLM. The significance of each coefficient was assessed using a Wald z-test. Significant coefficients (p < 0.01) were converted to their sign (±1), indicating whether higher spike counts during a given theta cycle significantly predicted leftward sweeps (+1, blue-colored cycles in Figure S7) or rightward sweeps (−1, red-colored cycles in Figure S7) at the bifurcation. Interestingly, at the p < 0.01 level as in the previous GLM analysis, we observed only a single cycle in which dSC turn cell activity predicted sweeps at the bifurcation, and this cycle involved activity from a left turn cell predicting rightward sweeps (Figure S7A). Implementing a less conservative significance threshold (p < 0.05), we did observe an increased proportion of cells and cycles in which dSC turn cell activity predicted future sweeps in a directionally consistent manner (Figure S7B); however, after performing permutation tests (1000 iterations, as described in the previous GLM analysis), we found that these proportions did not reliably exceed shuffled controls (Figure S7C).

### Quantifying deviations in behavioral trajectory around the turn

We used two approaches to compare animals’ behavioral trajectories around the bifurcation between ipsi and contra sweep trials. The first involved quantifying trajectory divergence from a linear path. To obtain a linear path, we first calculated the average trajectories during ipsi and contra sweep trials for turns in a given direction. We did so by interpolating each trial to the same number of samples and rotating them to a common coordinate frame (Figure 4A). We then defined a band of interest starting with an upper boundary near the bifurcation point of the maze and a lower boundary extending 10 cm from the upper boundary toward the base of the maze. This band was chosen to focus on trajectory deviations immediately prior to the bifurcation. The linear path was defined by connecting the average positions at which the ipsi and contra sweep trajectories crossed the upper and lower boundaries of the band, producing a single linear path against which trajectory deviations were measured. We then calculated, within this band of interest, the deviation of each ipsi and contra sweep trial from the linear path as the average perpendicular distance from each sample point to the line, producing a single deviation value per trial.

For the ROC analysis, deviation values for ipsi and contra sweep trials were treated as two distributions, and an ROC curve was computed. The optimal classification threshold was found by minimizing the distance to the top-left corner of the ROC curve. We calculated the balanced classification accuracy (% accuracy, (true positive rate + true negative rate) / 2) to assess ROC performance. For each session, classification accuracy was calculated separately for left and right turn trials (Figure 4B, left panel). For each animal, classification accuracies were averaged across turn directions and sessions (Figure 4B, right panel).

Trajectory deviations between ipsi and contra sweep trials were also quantified using the IdPhi measure (the integrated absolute angular velocity)^24^ calculated over the animal’s trajectory (Figure 4C). Briefly, average trajectories for ipsi and contra sweep trials were first smoothed (Gaussian, 1 sample) and heading angle was derived. IdPhi was defined as the sum of the absolute change in heading angle across each average trajectory, with higher values indicating greater deviations in heading. For each session, IdPhi was averaged across left and right turn trials, yielding one ipsi and one contra value per session (Figure 4C).

## Quantification and statistical analysis

All analyses and statistical tests were implemented using custom Python scripts (Python version 3.8.5). Statistical tests used and p-values are provided throughout the text and in figure legends.

## Supplemental Figures

**Figure S1:**
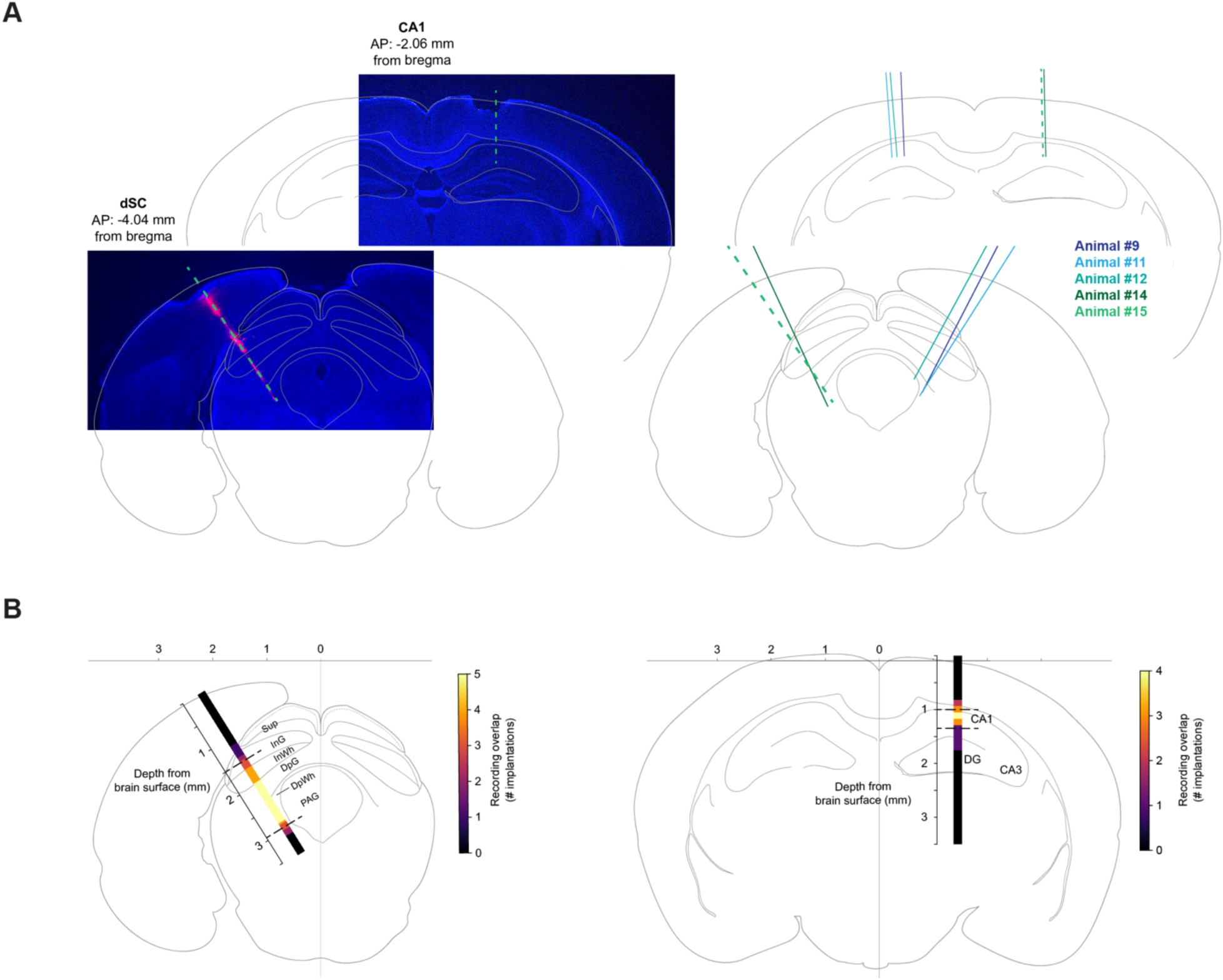
Histology and recording site analysis. **(A)** Left, example coronal sections of probe implantations targeting the dSC and dorsal CA1 regions in an example animal (animal #15). Probe track in dSC is marked by DiI (pink) and indicated by a dashed green line. Probe track in CA1 is indicated by a dashed green line. Right, probe tracks in dSC and dorsal CA1 across all implantations (n = 5 implantations from 5 mice). **(B)** Left, heatmap showing the approximate span of recording depths in the SC across implantations. Lighter colors indicate higher overlap. Dashed black lines indicate the average depths from the brain surface of the first and last dSC recordings. Note high concentration of recordings between the intermediate gray (InG) and deep white layers (DpWh). Heatmap is oriented at the average implantation angle and ML/AP coordinates across animals. (Sup = superficial layer, InG = intermediate gray layer, InW = intermediate white layer, DpG = deep gray layer, DpWh = deep white layer, PAG = periaqueductal gray). Right, same as left panel but for CA1 implantations. All outlines of histological sections were adapted from The Mouse Brain in Stereotaxic Coordinates^44^.

**Figure S2:**
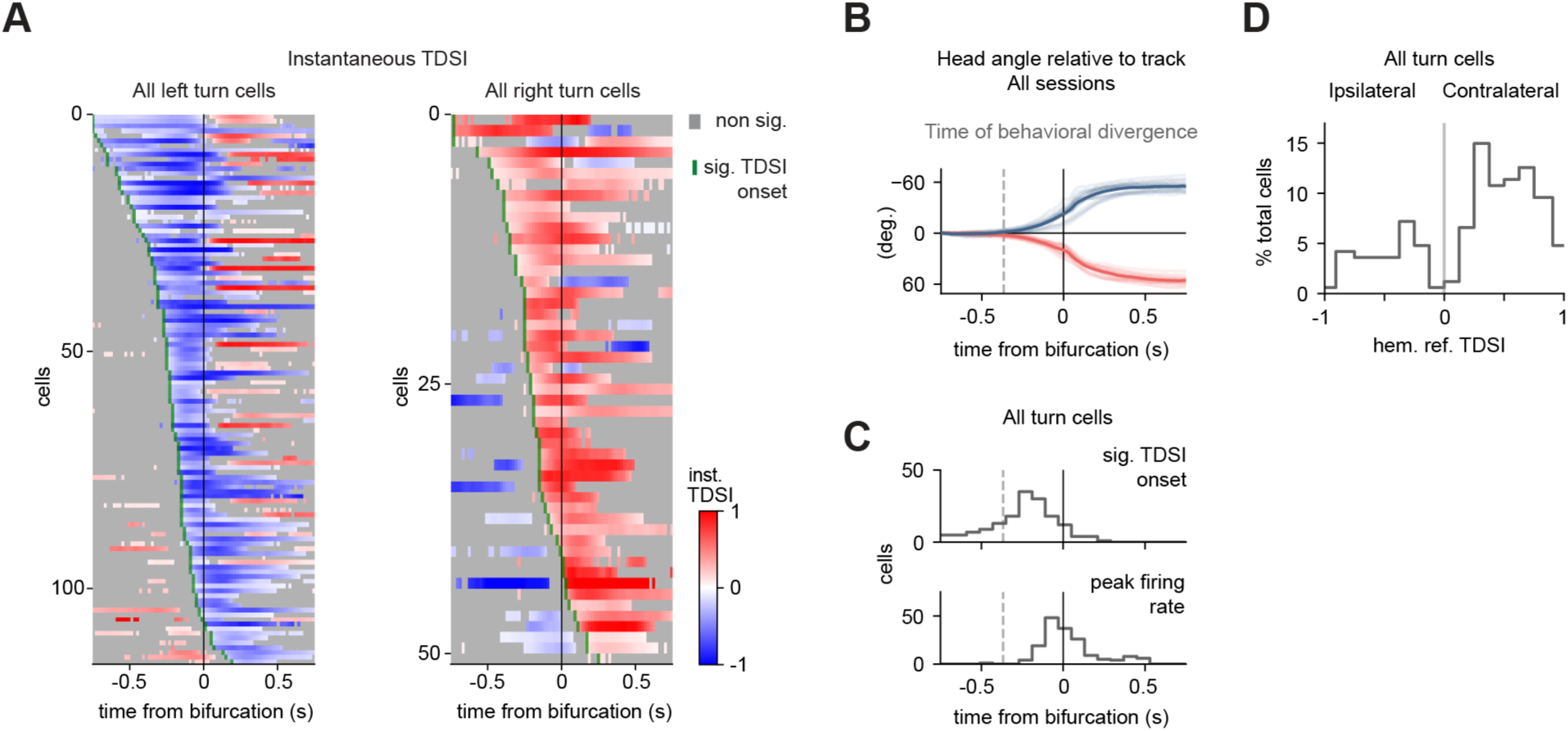
Characterization of dSC turn cells. **(A)** All left and right turn cells sorted by the time at which the instantaneous TDSI became selective in the cell’s preferred turn direction (significant TDSI onset; n = 116 left turn cells, n = 51 right turn cells; 3 right dSC insertions, 12 sessions; 2 left dSC insertions, 7 sessions). **(B)** Angle of the animals’ head relative to the track during the task. Dark lines: average over all sessions and animals (n = 19 sessions, 5 mice); light lines: individual sessions. Dashed line: time from the maze bifurcation at which the average head angle relative to the track significantly diverges between right and left turns (−0.37 s). **(C)** Distributions of significant TDSI onset times and times of peak firing rate across all turn cells. Dashed lines: average time of behavioral divergence as shown in (B). **(D)** Distribution of TDSI values across all turn cells referenced to the recorded dSC hemisphere. Positive and negative hemisphere referenced TDSI values indicate turn cells whose firing is selective for contraversive (e.g., left turn cells in the right dSC) or ipsiversive (e.g., right turn cell in right dSC) turns, respectively.

**Figure S3:**
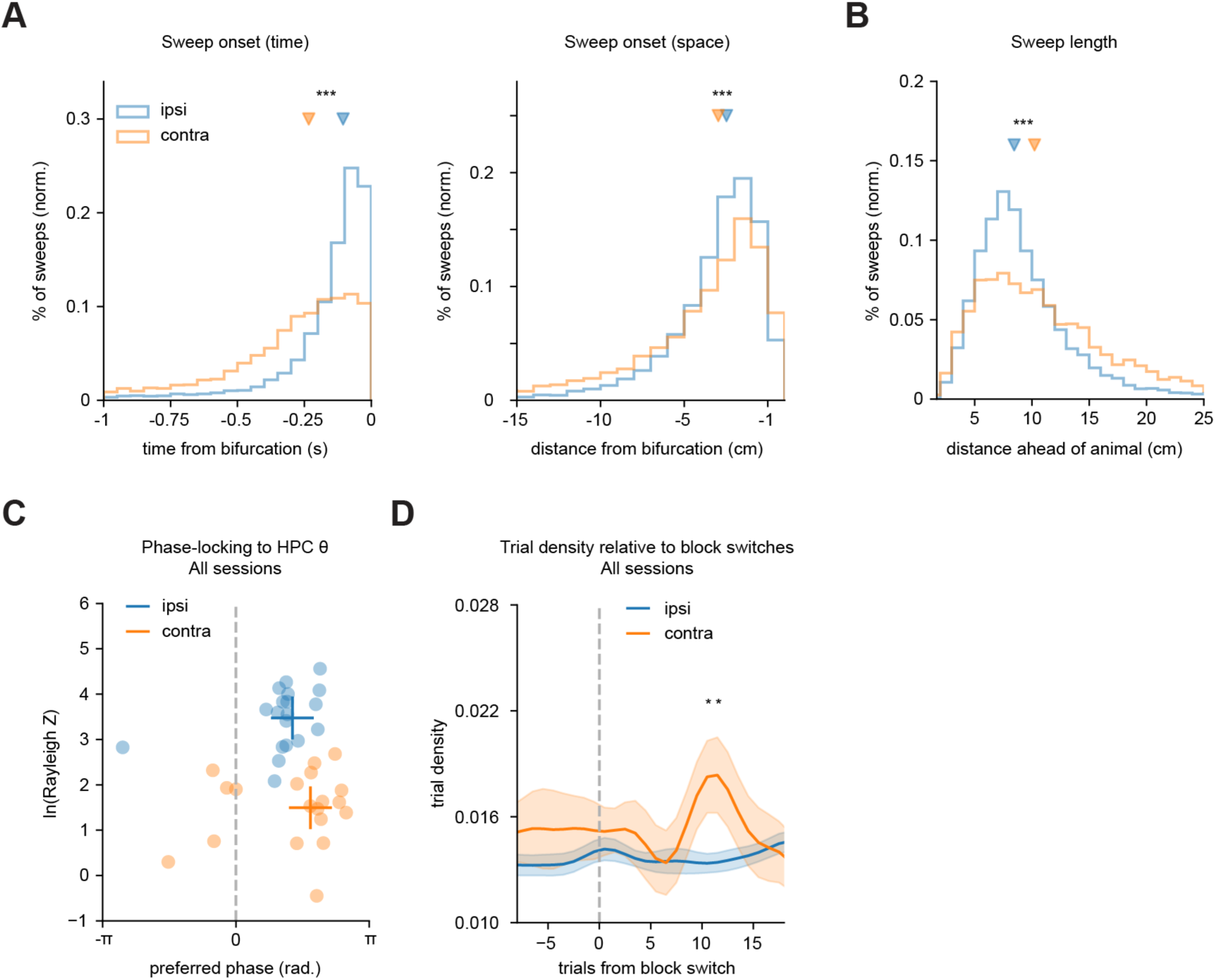
Ipsi contra sweep parameters. **(A)** Left, distributions of sweep onset times relative to the turn for ipsi and contra sweeps (ipsi sweeps, blue, n = 9178 sweeps, median: −105 ms, IQR: −201 to −54 ms, contra sweeps, orange, n = 3002 sweeps, median: −234 ms, IQR: −403 to −115 ms, p < 0.001, Mann-Whitney U test). Right, distributions of the animals’ distance from the turn at the onset of ipsi and contra sweeps (ipsi sweeps, median: −3.47 cm, IQR: −5.40 to −2.21 cm, contra sweeps, median: −3.96 cm, IQR: −7.31 to −2.20 cm, p < 0.001, Mann-Whitney U test). **(B)** Distributions of sweep distance from the animal’s position for ipsi and contra sweeps (ipsi sweeps, median: 8.45 cm, IQR: 6.47–11.29 cm, contra sweeps, median: 10.23 cm, IQR: 6.76–14.93 cm, p < 0.001, Mann-Whitney U test). Sweep length was calculated as the average difference between the animal’s actual and decoded positions during the duration of the sweep. **(C)** Phase-locking of sweeps to late phases (0 to π rad.) of the hippocampal theta cycle across sessions (n = 19 sessions, ipsi, pref. phase: +1.33 ± 0.13 rad., contra, pref. phase: +1.76 ± 0.26 rad., ipsi vs. contra: p > 0.05, permutation test). X-axis indicates theta phase where phase 0 corresponds to the trough of the theta cycle at the CA1 pyramidal cell layer. Y-axis indicates phase-locking strength (Rayleigh Z, see Methods). Note that, while both types of sweeps were phase-locked to late phases of theta, contra sweeps were more weakly phase-locked compared to ipsi sweeps (ipsi, ln(Z): 3.48 ± 0.15, contra, ln(Z): 1.49 ± 0.18, ipsi vs. contra: p < 0.001, paired t-test; the same effect was obtained using the mean resultant length). Cross-hair markers indicate averages. **(D)** Density of trials containing only ipsi or contra sweeps aligned to block switches across sessions (n = 19 sessions). Traces were normalized by the number of trials of each type (ipsi or contra) per session and smoothed with a Gaussian kernel (2 bins). Solid lines represent the mean and shaded bands represent s.e.m. Note the transient increase in the density of contra sweep trials following block switches relative to ipsi sweep trials (p < 0.05 per bin, permutation tests), consistent with recent findings in rats showing an increased proportion of representations of alternative possibilities during periods of increased cognitive demand^25^.

**Figure S4:**
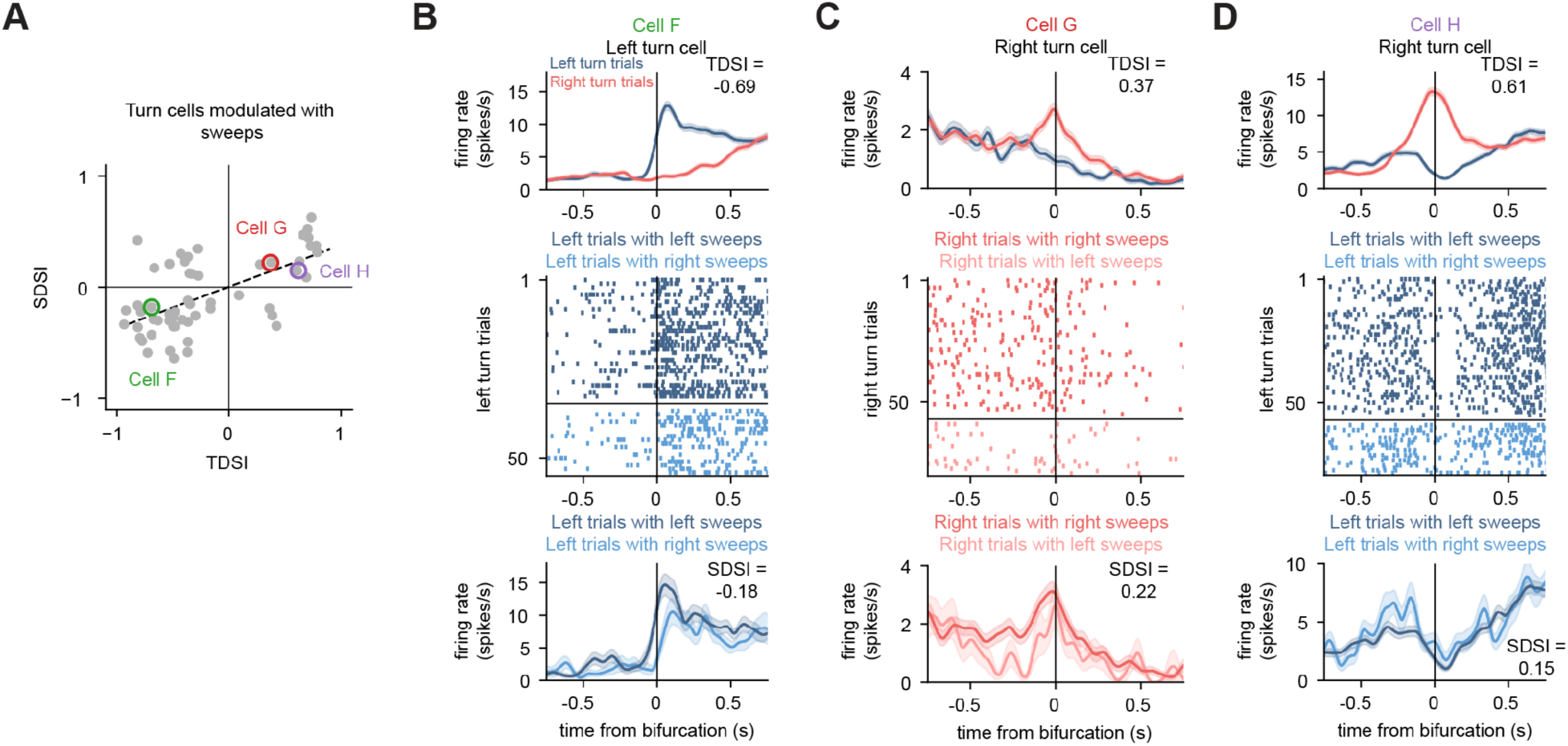
Additional examples of sweep-selective turn cells. **(A)** Same scatter plot as in Figure 2E. Markers indicate three additional sweep-selective turn cells (Cells F, G, and H) shown in panels B-D. **(B)** Example of left turn cell firing more during left turns with left sweeps compared to left turns with right sweeps (p < 0.05, permutation test). **(C)** Example of right turn cell firing more during right turns with right sweeps compared to right turns with left sweeps (p < 0.01, permutation test). **(D)** Example of right turn cell firing more during left turns with right sweeps compared to left turns with left sweeps (p < 0.05, permutation test).

**Figure S5:**
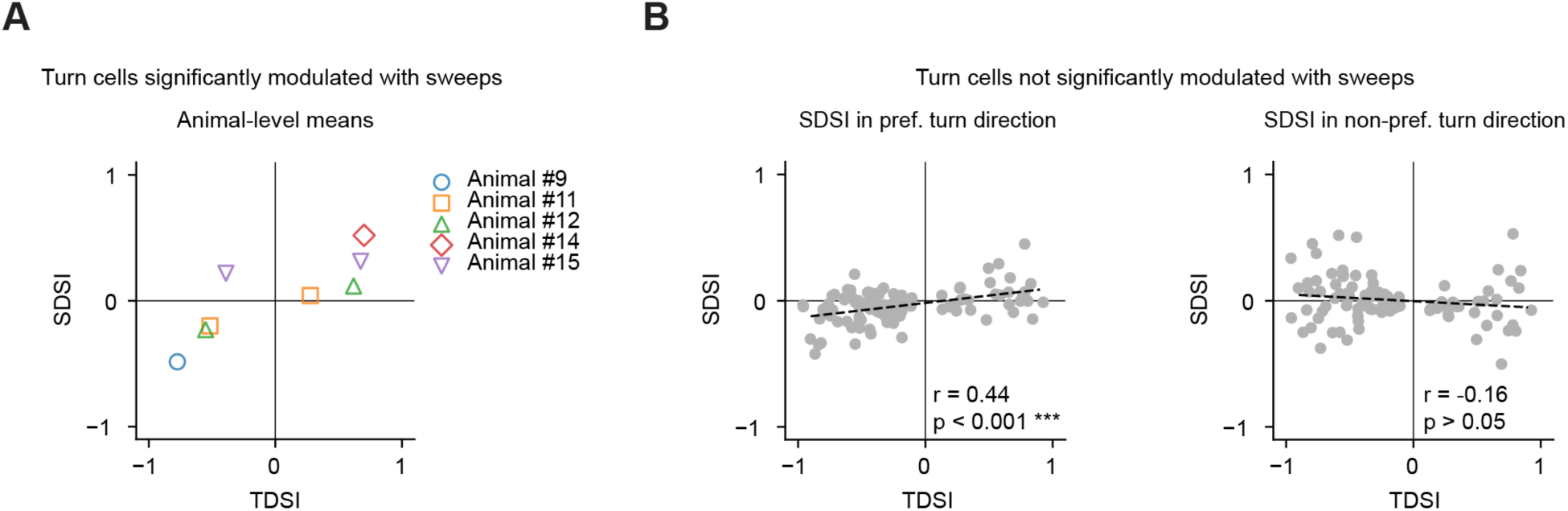
Relationship between TDSI and SDSI across animals and turn cells not significantly modulated with sweeps. **(A)** Correlation between TDSI and SDSI across individual animals. Animal-level means were calculated from cells whose activity was significantly modulated in coordination with sweep direction (Figure 2E). For each animal, cells were split by the sign of the TDSI (TDSI < 0 and TDSI > 0) and the mean TDSI and mean SDSI were computed for each half. Symbols show each animal’s mean TDSI vs. mean SDSI for each half (different symbol shapes and colors correspond to different animals, indicated to the right). Animals with cells in only one TDSI range are shown as a single point. **(B)** Left, correlation between TDSI and SDSI across cells whose activity was not significantly modulated in coordination with sweep direction (n = 98 cells). SDSI values in this panel were computed from trials in which the animal turned in each cell’s preferred turn direction (i.e., SDSI values for left turn cells during left turn trials, right turn cells during right turn trials, Pearson r = 0.44, p < 0.001). Right, same as left panel but with SDSI values computed from trials in which the animal turned in the cell’s non-preferred turn direction (i.e., SDSI values for left turn cells during right turn trials, right turn cells during left turn trials, Pearson r = −0.16, p > 0.05).

**Figure S6:**
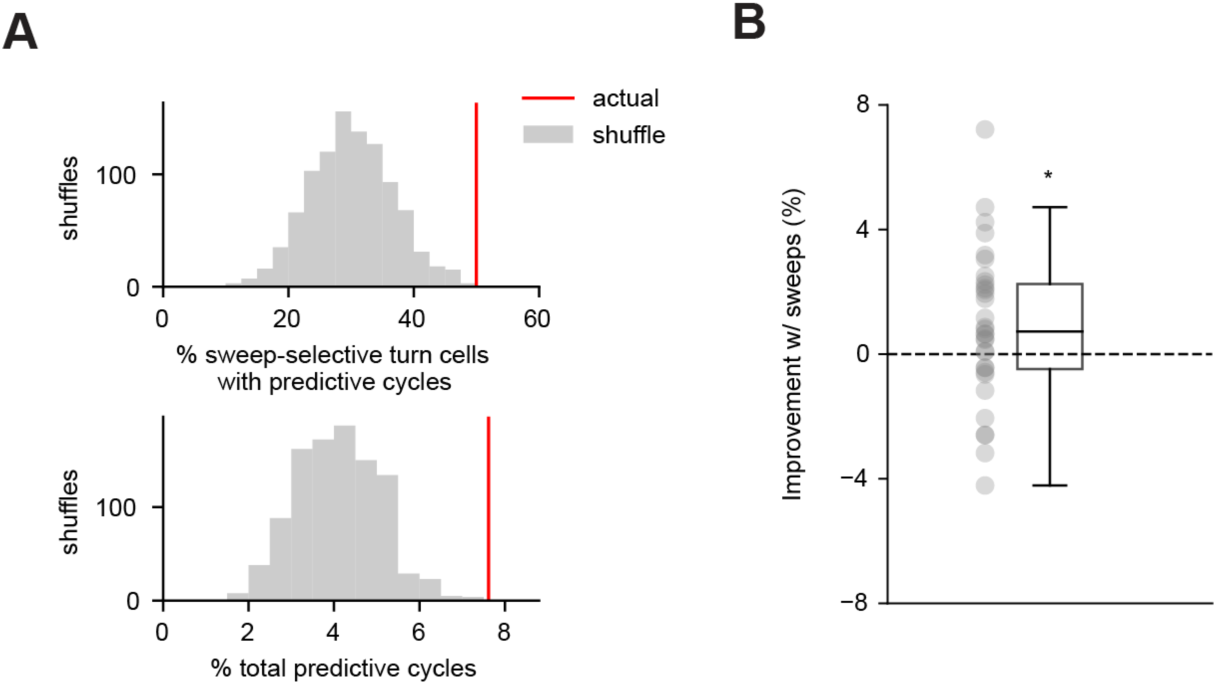
GLM shuffle controls and cross-validation for. Figure 3 **(A)** Top, distribution of the percentage of sweep-selective turn cells with predictive cycles after shuffling spike counts across trials to disrupt sweep-firing relationships (n = 1000 shuffles, shuffled: 29 ± 0.2%, actual = 50%, p = 0.001: 1 shuffle had a percentage of sweep-selective turn cells that exceeded that of the actual data). Bottom, same as top but for percentage of significantly predictive cycles (n = 1000 shuffles, shuffled: 4.1 ± 0.03%, actual = 7.6%, p < 0.001: 0 shuffles had a percentage of predictive cycles that exceeded that of the actual data). **(B)** Model improvement with sweep information compared to turn direction information alone for each significantly predictive cycle (n = 32 cycles from Figure 3B, mean: 0.89 ± 0.43% (s.e.m.), 95% CI [0.054, 1.72] %, p < 0.05, 1-sample t-test compared to 0% improvement). Model improvement is defined as percentage reduction in mean absolute error (MAE) when adding sweep predictors (full model) to a turn-only model (reduced model), calculated as: (MAE reduced model - MAE full model) / MAE reduced model × 100. Positive values indicate model improvement with sweeps. For cross-validation, values were calculated and averaged across folds of a 5-fold train-test split.

**Figure S7:**
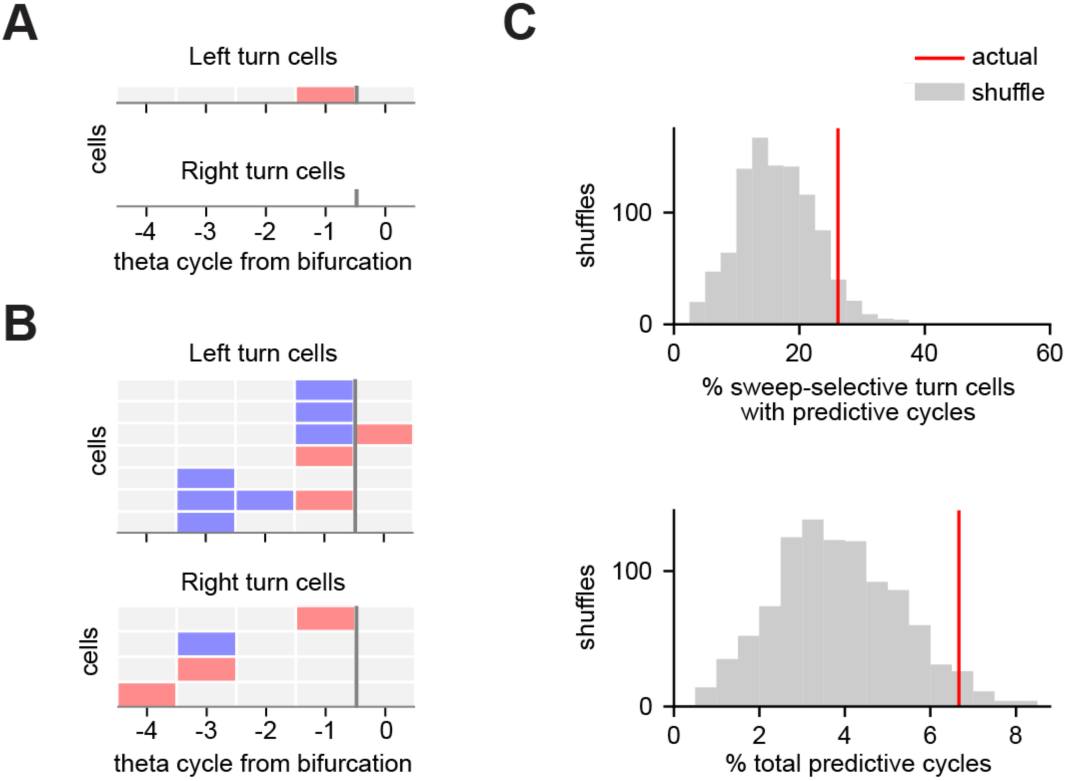
Turn cell activity does not reliably predict future hippocampal sweeps. **(A)** GLM analysis to assess whether dSC turn cell activity during individual theta cycles prior to and at the time of the bifurcation (cycles −4 through −1 and cycle 0, respectively) can predict the direction of hippocampal sweeps at the bifurcation (see Methods). Shown are all cells with predictive cycles at a p < 0.01 threshold (Wald z-tests). Shaded red boxes indicate cycles during which dSC firing predicted rightward sweeps at cycle 0. Shaded blue boxes, if present, indicate cycles during which dSC firing predicted leftward sweeps at cycle 0. Note that only activity for one left turn cell at cycle −1 predicted rightward sweeps at cycle 0, and no right turn cells exhibited predictive cycles. **(B)** All left and right turn cells with predictive cycles at a p < 0.05 threshold (Wald z-tests) sorted by theta cycle relative to the bifurcation. Color code is same as in A. Note increased number of cells with predictive cycles (i.e., theta cycles during which dSC activity significantly predicted sweeps at the bifurcation) compared to A. **(C)** Shuffle controls for cells and cycles in B (p < 0.05 threshold). Top, distribution of the percentage of sweep-selective turn cells with predictive cycles after shuffling spike counts across trials to disrupt firing-sweep relationships (n = 1000 shuffles, shuffled: 17 ± 0.2%, actual = 26%, p = 0.079: 79 out of 1000 shuffles had a percentage of sweep-selective turn cells that exceeded that of the actual data). Bottom, same as top but for percentage of significantly predictive cycles (n = 1000 shuffles, shuffled: 3.9 ± 0.04%, actual = 6.7%, p = 0.047).

**Figure S8:**
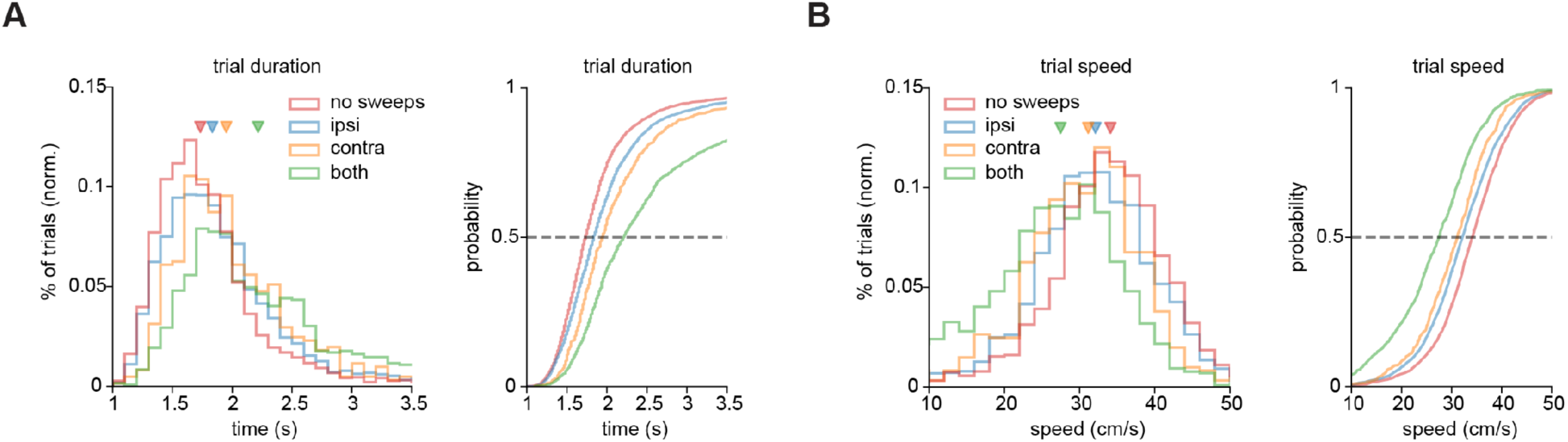
Effects of sweep presence and direction on trial duration and speed. **(A)** Left, trial durations for trials with no sweeps (n = 2810 trials, red), ipsi sweeps (n = 3944, blue), contra sweeps (n = 607, orange), and both ipsi and contra sweeps (n = 1290, green) pooled across sessions and animals (n = 19 sessions, 5 animals). Markers indicate medians of each group (1.73, 1.83, 1.95, and 2.22 s; IQRs: 1.52–2.00, 1.58–2.18, 1.68–2.35, 1.82–2.95 s, respectively, p < 0.001 for all comparisons, Mann-Whitney U tests, Bonferroni-corrected). Right, cumulative plot of each distribution in the left panel. **(B)** Left, trial speeds for trials with no sweeps, ipsi sweeps, contra sweeps, and both ipsi and contra sweeps pooled across sessions and animals as in A. Speed was calculated as a single value across each trial as 2D distance traveled divided by trial duration. Markers indicate medians of each group (34.1, 32.1, 31.2, 27.5 cm/s; IQRs: 29.6–38.9, 27.3–37.2, 26.0–35.4, 21.3–32.7 cm/s, respectively, p < 0.001 for all comparisons, Mann-Whitney U tests, Bonferroni-corrected). Right, cumulative plot of each distribution in the left panel.

